# Multiple mechanisms of self-association of chemokine receptors CXCR4 and CCR5 demonstrated by deep mutagenesis

**DOI:** 10.1101/2023.03.25.534231

**Authors:** Kevin S. Gill, Kritika Mehta, Jeremiah D. Heredia, Vishnu V. Krishnamurthy, Kai Zhang, Erik Procko

**Affiliations:** Department of Biochemistry, University of Illinois, Urbana, IL 61801, USA; Codexis, Redwood City, CA 94063; Cyrus Biotechnology, Seattle, WA 98121, USA

## Abstract

Chemokine receptors are members of the rhodopsin-like class A GPCRs whose signaling through G proteins drives the directional movement of cells in response to a chemokine gradient. Chemokine receptors CXCR4 and CCR5 have been extensively studied due to their roles in white blood cell development and inflammation and their status as coreceptors for HIV-1 infection, among other functions. Both receptors form dimers or oligomers but the function/s of self-associations are unclear. While CXCR4 has been crystallized in a dimeric arrangement, available atomic resolution structures of CCR5 are monomeric. To investigate the dimerization interfaces of these chemokine receptors, we used a bimolecular fluorescence complementation (BiFC)-based screen and deep mutational scanning to find mutations that modify receptor self-association. Many disruptive mutations promoted self-associations nonspecifically, suggesting they aggregated in the membrane. A mutationally intolerant region was found on CXCR4 that matched the crystallographic dimer interface, supporting this dimeric arrangement in living cells. A mutationally intolerant region was also observed on the surface of CCR5 by transmembrane helices 3 and 4. Mutations from the deep mutational scan that reduce BiFC were validated and were localized in the transmembrane domains as well as the C-terminal cytoplasmic tails where they reduced lipid microdomain localization. The reduced self-association mutants of CXCR4 had increased binding to the ligand CXCL12 but diminished calcium signaling. There was no change in syncytia formation with cells expressing HIV-1 Env. The data highlight that multiple mechanisms are involved in self-association of chemokine receptor chains.

## INTRODUCTION

G-protein coupled receptors (GPCRs) are integral membrane proteins that activate heterotrimeric G protein signaling pathways, leading to diverse downstream signals. They are involved in sight, hormonal regulation, white blood cell trafficking, and a wide range of other physiological functions. While it is accepted that many GPCRs dimerize, the functional consequences of dimerization on ligand interactions and downstream signaling remain incompletely understood, with structural experiments of class A rhodopsin-like GPCRs suggesting they bind to ligand as monomers (Kufareva et al., 2014), and coimmunoprecipitation and FRET experiments suggesting they are active as dimers (Albizu et al., 2010; Hebert et al., 1996; Kobayashi et al., 2017). Ligand-receptor interactions may have additional complexities where the ligands may act as monomers or dimers with different effects (Drurya et al., 2011) and there may be negative cooperativity between subunits in a GPCR dimer such that effectively only one of the receptors is ligand-bound (El-Asmar et al., 2005). To determine whether dimerization is important for GPCR activity, researchers have tried different methods to identify dimerization interfaces for class A GPCRs with limited success. While class C GPCRs such as GABA_B_ receptors, metabotropic glutamate receptors and sweet and umami taste receptors are obligate dimers (El Moustaine et al., 2012; Jones et al., 1998), the dimerization of class A GPCRs is less understood, although it has been observed in multiple crystal structures including those of rhodopsin (Zhao et al., 2019) and μ-opioid receptor (Manglik et al., 2012). Different transmembrane helices (TM) have been observed at dimer interfaces, including TM1 and TM4-6.

We have previously used conformation-dependent antibodies against the class A GPCRs CXCR4 and CCR5 to develop a method for deep mutational scanning (DMS) of transmembrane proteins expressed on human cells (Heredia et al., 2018). CXCR4 and CCR5 are chemokine receptors, which in response to a chemokine gradient cause cell migration, often of white blood cells, but also of other cell types and some cancers (Gassenmaier et al., 2013; Müller et al., 2001). These two chemokine receptors are also well studied for their role as coreceptors for HIV-1 infection (Connor et al., 1997; Deng et al., 1996; Dragic et al., 1996). There is extensive evidence for CXCR4 dimerization and oligomerization (Babcock et al., 2003; Lao et al., 2017; Martínez-Muñoz et al., 2018; Vila-Coro et al., 1999) and the protein crystallizes as a dimer (Wu et al., 2010). Wu et al. showed a dimerization interface using TM3-5 and the extracellular tip of TM6 that was consistent across 5 different crystal forms. Structures of CCR5 have been determined with and without ligands as a monomer (Peng et al., 2018; Tan et al., 2013; H. Zhang et al., 2021) but not yet as a homodimer. Models for CCR5 dimerization have been made through comparisons to other class A GPCR dimeric structures and molecular dynamics simulations (Jin et al., 2018; F. Zhang et al., 2019). These methods have suggested multiple different dimerization interfaces that collectively cover almost all sides of the protein. Given GPCRs are known to oligomerize and hetero-oligomerize with other GPCRs (Fonseca & Lambert, 2009; Sohy et al., 2009), finding physiologically relevant dimerization interfaces has been challenging.

By examining the mutational landscapes of CXCR4 and CCR5 for binding to antibodies 12G5 and 2D7, respectively, we found conserved sites on the surface of the transmembrane domains that, with respect to CXCR4, matched previous crystallographic dimers (Wu et al., 2010). To further elucidate how these chemokine receptors dimerize, we used bimolecular fluorescence complementation (BiFC) with deep mutational scanning to search for membrane-exposed surface patches that were intolerant of mutations for high BiFC signal. We found regions on the chemokine receptors that are hypothesized sites for dimerization, which on CXCR4 again matched closely with the crystallographic dimer conformation. We also found many structurally-disruptive mutations that increased BiFC signal, including nonspecifically with an unrelated GPCR, suggesting overexpression of mutant chemokine receptors may lead to aggregation. Finally, mutations were found in the cytoplasmic tails that reduced BiFC between receptors and also reduced colocalization with a marker for lipid microdomains. We conclude that there are multiple mechanisms that mediate self-association of chemokine receptor polypeptides.

## EXPERIMENTAL PROCEDURES

### Cell Culture

The X4-KO Expi293F and the MA-VN expressing X4-KO Expi293F cell lines are previously described (Heredia et al., 2018, 2019). Cells (Thermo Fisher) were cultured in Expi293 Expression Media (Thermo Fisher) at 37 °C, 8% CO_2_, and 125 rpm. Cells were transfected at a density of 2 × 10^6^ cells/ml with 500 ng plasmid DNA per ml of culture using Expifectamine (Thermo Fisher) according to the manufacturer’s directions. To generate stable lines for BiFC, X4-KO Expi293F cells were transfected with linearized pCEP4-FLAG-CXCR4-VN or pCEP4-FLAG-CCR5-VN and selected with hygromycin B (200 μg/ml). The population of CXCR4-VN or CCR5-VN positive cells was enriched by FACS, in which cells were washed in Dulbecco’s phosphate buffered saline (PBS) supplemented with 0.2% bovine serum albumin (BSA), stained with 1:200 chicken anti-FLAG-FITC (Immunology Consultants Laboratory) in PBS-BSA, washed and resuspended for sorting on a BA FACS Aria II. The collected cells were expanded by culturing in Expi293 Expression Medium supplemented with penicillin-streptomycin.

### Plasmids

Plasmids for myc-tagged CCR5 and CXCR4, used in signaling and syncytia assays, are previously described (Heredia et al., 2018) and available on Addgene (# 98948 and # 98946). Plasmids for myc-CXCR4-VC (Addgene # 98967), myc-CCR5-VC (# 98966), myc-GRM3-VC (# 98968), FLAG-CXCR4-VN (# 98964), FLAG-CCR5-VN (# 98963), and FLAG-GRM3-VN (# 98965) are previously described (Heredia et al., 2018). Briefly, the N-terminal half (VN) of the yellow fluorescent protein Venus (a.a. 1-154; mutant I152L) or the C-terminal half (VC) of Venus (a.a. 155-238) were fused to the C-termini of tagged receptors and cloned into the NheI-XhoI sites of pCEP4 (Invitrogen). Plasmids for HIV-1 MN gp160 and BaL gp160 (# 100919) are previously described (Heredia et al., 2018). pCMV3-CD4 was from Sino Biological (# HG10400-UT). For single molecule imaging, a halo tag sequence was inserted into the myc-CXCR4 wild type and low BiFC mutant sequences, in between the myc tag and the chemokine receptor sequence.

### Library Generation

Single site-saturation mutagenesis (SSM) libraries of human myc-tagged CXCR4 and CCR5 were generated previously (Heredia et al 2018). Here, the libraries were modified by using PCR-based fragment assembly to fuse the 3’ ends of mutated chemokine receptor sequences to VC (a.a. 155-238 of Venus). The PCR products were digested and ligated into the NheI-XhoI sites of pCEP4 and electroporated into *E. coli* DH5α cells. The number of transformants was > 100x the theoretical sequence diversity. The myc-CXCR4-VC and myc-CCR5-VC plasmid libraries were transfected into stable FLAG-CXCR4-VN or FLAG-CCR5-VN Expi293F lines under conditions where cells typically express no more than one mutant VC-fused chemokine receptor gene per cell: 1 ml culture at a density of 2 × 10^6^ cells was transfected with 1 ng library DNA diluted with 1.5 μg pCEP4ΔCMV carrier DNA using Expifectamine (Thermo Fisher). The media was replaced 2 h post-transfection and cells were harvested for sorting 24 h post-transfection.

### Sorting Cells for Surface Expression and High BiFC Signal

Cells transfected with the libraries were washed with cold PBS-BSA and incubated with 1/300 anti-myc Alexa 647 (clone 9B11; Cell Signaling Technology) plus 1/300 anti-Flag Cy3 (clone M2; Sigma-Aldrich) in PBS-BSA for 30 minutes before being washed twice and resuspended in cold PBS-BSA supplemented with 1/100 fetal bovine serum (FBS). Cells were sorted on a BD FACS Aria II at the Roy J. Carver Biotechnology Center. Cells were first gated by forward-side scatter and side scatter for the main population of cells. For sorting based on surface expression, cells positive for the myc tag (Alexa 647 fluorescence) were collected. For sorting based on high BiFC signal, within the expression gate the top 5% of cells for YFP/Venus fluorescence relative to surface receptor expression were collected. Cells were sorted for no longer than 4 h to maintain high viability and were collected in tubes coated overnight in FBS. Collected cells were centrifuged and the pellets stored at -80 °C.

### Illumina Sequencing and Analysis

RNA was extracted from frozen cells from FACS using the GeneJET RNA Purification Kit (Thermo Fisher) and reverse transcribed with AccuScript primed with the EBV reverse sequencing primer for first strand cDNA synthesis. The receptor cDNA was PCR amplified as three overlapping fragments to achieve full coverage of the gene. In a second round of PCR, experiment-specific barcodes and adaptors for annealing to the Illumina flow cell were added. Samples were sequenced (2 × 250 nt) using Illumina HiSeq 2500 at the UIUC Roy J. Carver Biotechnology Center. Data were analyzed using Enrich (Fowler et al., 2011). Log_2_ enrichment ratios of mutants were adjusted by subtracting the enrichment of the wild type sequence. Data and analysis commands are deposited with NCBI’s Gene Expression Omnibus under series accession number GSE125426.

### Validation of Receptor Mutants with Changes in BiFC

pCEP4 plasmids containing wild type VN and VC fused receptors were used as templates for targeted mutagenesis by overlap extension PCR. Mutated inserts in all plasmids were confirmed by Sanger sequencing. Plasmids were transiently transfected as described above into the relevant cell line and analyzed by flow cytometry for receptor expression and BiFC signal 24 h post-transfection. Cells were washed with PBS-BSA, incubated in 1:100 anti-myc Alexa 647 and 1:100 anti-FLAG Cy3 in PBS-BSA for 20 minutes, washed twice, and resuspended in PBS-BSA for analysis on a BD LSR II. Data were collected using instrument software and analyzed with FCS Express (De Novo Software). Cells were gated by forward-side scatter for the main population, and then gated by Alexa 647/Cy3 fluorescence to control for a consistent level of surface expressed receptor across the samples. Mean YFP/Venus fluorescence was recorded.

### Co-Localization with Matrix

Using a MA-VN cell line (Heredia et al., 2019), cells were transiently transfected as described above with wild-type or mutant myc-tagged receptors fused to VC and cloned into pCEP4. Cells were processed 24 h post-transfection. Cells were washed PBS-BSA and incubated in a 1:200 dilution of anti-myc Alexa 647 (clone 9B11; Cell Signaling Technology). Cells were washed and analyzed on a BD LSRII flow cytometer.

### CXCL12 Binding Assay

Transfected X4-KO Expi293F cells were harvested 24 h post-transfection, washed with PBS-BSA and incubated in 1:200 anti-myc-Alexa 647 (clone 9B11; Cell Signaling Technology) and 10 μM CXCL12-sfGFP for 30 minutes. Cells were washed and analyzed on a BD LSR II. The preparation of CXCL12-sfGFP is described elsewhere (Heredia et al., 2018).

### Calcium Mobilization

X4-KO Expi293F cells were transiently transfected as described above with myc-tagged CXCR4 or CCR5 plasmids. Cells were prepared at room temperature 24 h post-transfection. Cells were washed with assay buffer (PBS containing 0.2% BSA and 1 mM CaCl_2_) and resuspended in assay buffer with 2 μM Fluo-4-AM (Life Technologies). Fluo-4-AM was prepared as a 1 mM stock in DMSO. Cells were incubated with 1:250 anti-myc-Alexa-647 (clone 9B11; Cell Signaling Technology) for 30 minutes with frequent mixing, and washed and resuspended in assay buffer. Cells were analyzed on a BD Accuri C6 Cytometer. Cells were gated by forward-side scatter for the main population and the Fluo-4 fluorescence of the Alexa 647 positive population was monitored over time. To cells at baseline, CXCL12 (PeproTech) or CCL5 (PeproTech) were added at final concentrations of 1, 10 and 100 ng/mL. Fluorescence returned to baseline within 120 s, at which point ionomycin was added to a final concentration of 4 μM. The spike in Ca2+-dependent Fluo-4 fluorescence in response to the chemokine was normalized to the maximal response induced by ionomycin.

### Syncytia Formation Assay

X4-KO Expi293F cells (2 × 10^6^ cells in 1 ml) were transfected with 50 ng pCMV3-CD4 and 450 ng pCEP4-myc-CXCR4 or pCEP4-myc-CCR5. A separate set of cells (2 × 10^6^ cells in 1 ml) were transfected with 500 ng pCEP4-gp160 from the MN or BaL HIV-1 strains. Empty vector was used for control transfections. After 5 h incubation at 37 °C, 8% CO_2_, 125 rpm, 0.2 × 10^6^ receptor-expressing cells and gp160-expressing cells were mixed and added to 0.6 ml Expi293 Expression Medium. The cells were added to wells of a 12-well tray that had been incubated for 30 minutes with 0.01% poly-L-lysine (Sigma) to coat the plastic surface. Cells were incubated without agitation for 20 h. Wells were washed with warm PBS and attached cells removed with 0.25% trypsin-2.21 mM EDTA for 15 minutes at 37 °C. Detached cells were washed with cold PBS-BSA and analyzed on a BD LSR II flow cytometer to detect syncytia based on high forward-side scattering. The positive gate was set at less than 1% of negative control cells.

### Single Molecule Imaging of CXCR4

Coverslips measuring 24 mm × 40 mm (VWR Cat. No. 48393230) were cleaned for single molecule experiments as described previously (Camp et al., 2020). The washed coverslips were stored in molecular grade water, air-dried, and flamed for a few seconds. Coverslips were then immersed in a clean petri dish with 50 μg/mL poly-L-Lysine (PLL; Cat. No. P1274; Sigma-Aldrich) and incubated overnight. Coverslips were rinsed twice in molecular grade water and air dried. A large PDMS chamber was carefully assembled over a dried coverslip and placed in a 60 mm dish.

X4-KO Expi293F cells were transfected with wildtype or mutant myc-halo-CXCR4. 2 h post-transfection, cells were diluted 1:10 in Expi293 Expression Media and placed on the assembled PDMS coverslip chamber. 24 h post-transfection, cells were gently washed on the coverslip with PBS containing calcium and magnesium (Cat. No. 21-030CM; Corning) and incubated in 2 nM JF549-Halo-ligand (Cat. No. GA1110; Promega) for 15 minutes at 37 °C. Cells were washed three more times and incubated in PBS for another 15 minutes before four more washes and covered in PBS.

Single molecule tracking was performed at room temperature in a custom-built TIRF microscope (TIRFM). A 100X oil immersion objective (100X, N.A. 1.49, oil immersion) was assembled on an inverted microscope. A 561 nm laser was used to excite the labelled molecules. Power of the laser was controlled using neutral density filters. Individual labeled molecules were seen as diffraction-limited spots on the cell surface in immediate contact with the coverslip. The fluorescence from the single spots was collected by the same objective, passing an emitter and captured by an Electron Multiplying Charge Coupled Device (EMCCD) camera. A total of 2400 frames/trajectory were acquired for each field-of-view with an integration time of 50 ms. The collected data was exported to FIJI and single spots were tracked using the plugin TrackMate (Tinevez et al., 2017). Individual spots were selected as 3×3 pixel bright features. Threshold was selected such that all the single spots were selected in a frame. For track generation, the linking distance was fixed to 2 pixels, merging was not allowed, Gap closing distance was set as 2 pixels, and the Max frame gap was 2 frames. Data files were exported to MATLAB 2016b (MATLAB Release 2016b, The MathWorks, Inc.). For Divide and Conquer Moment Scaling spectrum transient diffusion analysis (DC-MSS), we used the MATLAB based software (Vega et al., 2018). All the tracks were segregated into super diffusion, free, confined, and immobile motions using DC-MSS.

## RESULTS

### Mutations to membrane-exposed surfaces of CXCR4 and CCR5 are deleterious for binding conformation-dependent antibodies

We revisited published deep mutational scans of the chemokine receptors CXCR4 and CCR5 to search for evidence of receptor oligomers (Heredia et al., 2018). These scans determined how mutations in the receptors impacted surface expression and recognition by conformation-dependent monoclonal antibodies: 12G5 for CXCR4 and 2D7 for CCR5. These antibodies bind to extracellular loops of the receptors, and we hypothesized that mutations in transmembrane regions that disrupt oligomeric organization may either alter surface expression or how the extracellular epitope is presented. Mapping conservation scores from the mutational scan of CXCR4 for binding to 12G5 onto the CXCR4 crystal structure (Wu et al., 2010) revealed that the dimer interface is indeed weakly but discernably more conserved than other membrane-exposed surfaces (**Figure 1A**). We thus turned our attention to CCR5, for which a dimeric structure has not been determined at atomic resolution. Mapping conservation scores from the mutational scan of CCR5 for binding to 2D7 to the CCR5 monomer structure (Tan et al., 2013), we found that a surface primarily formed by TM4 was more conserved than other membrane-exposed surfaces and might therefore form the dimer interface (**Figure 1B**). We thus introduced mutations into this region of CCR5 that were depleted in the scan for 2D7 binding (**Figure 1C**). Two control mutations located elsewhere on CCR5 and expected to have no effect were also evaluated. We found as expected that most of the mutations did indeed decrease 2D7 affinity (**Figure 1D**). To understand whether the mutations decreased receptor associations as hypothesized, we co-expressed CCR5 mutants fused to N-terminal (VN) or C-terminal (VC) segments of split Venus (a variant of yellow fluorescent protein or YFP). If the receptors closely associate, the VN and VC polypeptides will fold together to produce fluorescent Venus. This method is known as bimolecular fluorescence complementation (BiFC) (Ohashi & Mizuno, 2014). However, to our surprise, mutations that reduced CCR5 affinity for 2D7 caused increased BiFC signal (**Figure 1D**). This suggested that the mutations to the hypothesized dimer interface of CCR5 were generally disruptive of structure, reducing 2D7 recognition and causing putative aggregation in the membrane, measured as increased BiFC. These initial observations inspired us to use BiFC as the basis for deep mutational scans of CXCR4 and CCR5 to see whether we could identify mutations that decreased receptor associations.

**Figure 1.**
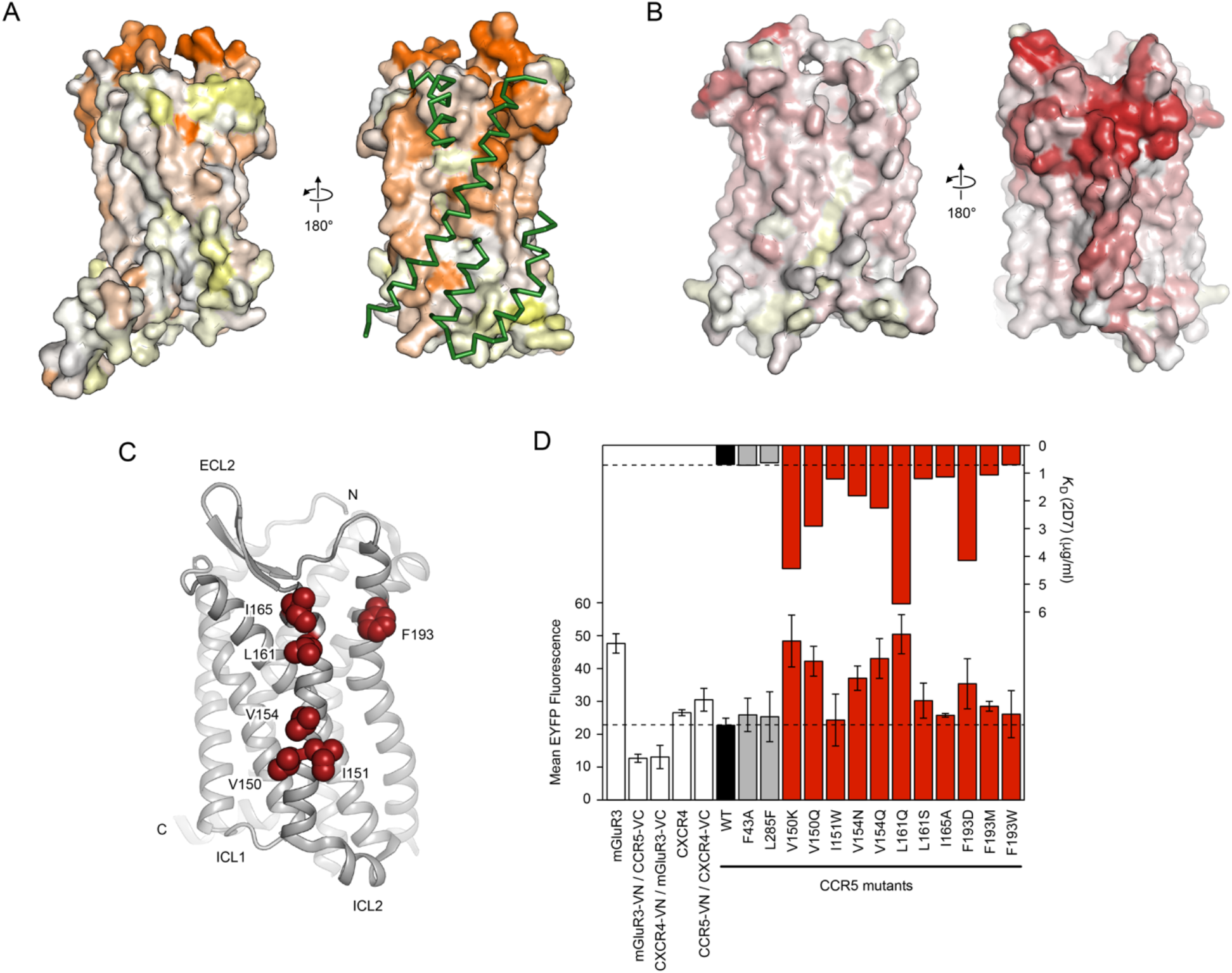
Deep mutational scans of chemokine receptors based on recognition by conformation-dependent antibodies highlights conserved surfaces facing the membrane. **(A)** Using the published deep mutational scan of CXCR4 for binding to monoclonal 12G5, experimental conservation scores are mapped to the CXCR4 surface (PDB 3ODU). Mutational tolerance of residues is shown from low in orange to high in white and yellow. At right, a more conserved surface exposed to the membrane corresponds to the interface for a second CXCR4 subunit in the crystal structure (regions in close contact are shown as green ribbons). **(B)** Conservation scores from the published deep mutational scan of CCR5 for binding to monoclonal 2D7 are mapped to the CCR5 crystal structure (PDB 4MBS), from conserved residues in red to mutationally tolerant residues in white and yellow. At right, one side of CCR5 exposed to the membrane is more conserved. **(C)** Cartoon of the CCR5 structure oriented with the conserved surface facing the reader. Residues targeted for mutagenesis are shown as red spheres. **(D)** *(Upper plot)* CCR5 mutants were expressed on Expi293F cells and the apparent K_D_ of 2D7 was measured by flow cytometry. Wild-type CCR5 and two control mutations not expected to impact 2D7 affinity are shown in black and grey, respectively. Mutations in the conserved membrane-exposed surface are shown in red. *(Lower plot)* Mutations were introduced into CCR5-VC and CCR5-VN constructs and BiFC measured in transfected Expi293F cells. Various controls are in white (see description in main text). Data are mean ± SD, n=3 independent replicates.

### Deep mutational scans of CXCR4 and CCR5 based on BiFC

Deep mutational scanning couples an *in vitro* screen or selection of a diverse library of sequence variants with next generation sequencing. BiFC is well suited to illuminating how mutations in chemokine receptors influence receptor self-association, as BiFC facilitates the separation of cells using fluorescence-activated cell sorting (FACS). Using the split Venus BiFC method (Ohashi & Mizuno, 2014), single site saturation mutagenesis (SSM) libraries of CXCR4 and CCR5 (Heredia et al., 2018) were modified by fusing the C-terminal half of split Venus to the C-termini, creating CXCR4-VC and CCR5-VC libraries. These were expressed in an Expi293F cell line that was modified to (i) first remove endogenous CXCR4 using CRISPR-Cas9 (CXCR4-knockout or X4-KO (Heredia et al., 2018)) and (ii) second to stably integrate a gene encoding either CXCR4 or CCR5 fused at their C-termini to the N-terminal half of split Venus, creating CXCR4-VN and CCR5-VN stable lines without endogenous CXCR4 expression. The VC and VN fused receptors had different N-terminal (and thus extracellular) epitope tags for their detection. The stable lines were transiently transfected with the matching VC-fused receptor libraries under conditions where the cells typically express no more than one gene per cell, providing a tight link from genotype to phenotype. The cell libraries were sorted by fluorescence-activated cell sorting (FACS), in which collection gates were applied to cells that expressed CXCR4-VC or CCR5-VC mutants at the plasma membrane. In a second sorting experiment, cells were gated not only for surface CXCR4-VC or CCR5-VC expression but also for high levels of Venus fluorescence (**Figure 2**), indicative of an association between a VC-fused mutant with a VN-fused wild-type receptor. RNA transcripts from the sorted libraries were analyzed by Illumina sequencing and compared to the naïve plasmid libraries to calculate an enrichment ratio for each mutation. The enrichment or depletion of single mutations in the CXCR4-VC and CCR5-VC genes are presented as heat maps that represent mutational landscapes for receptor associations **(Figures 3 and 4)**.

**Figure 2.**
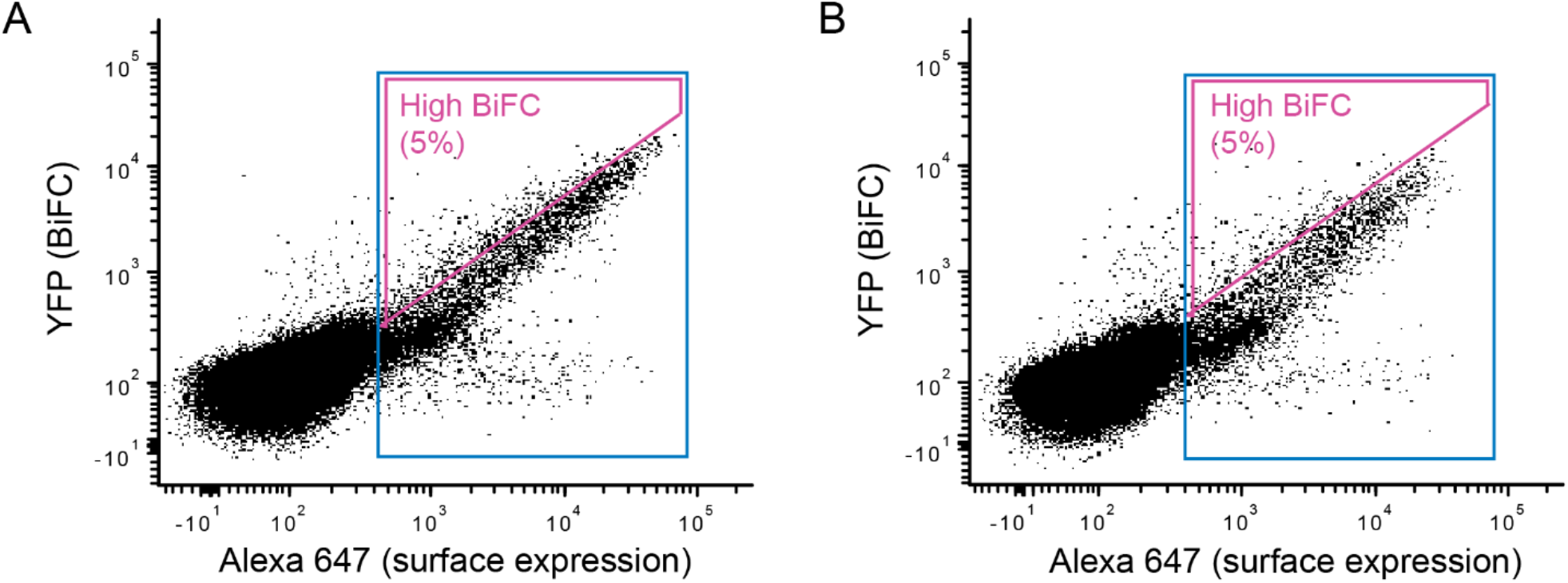
FACS collection gates for sorting human cells expressing SSM libraries of chemokine receptors CXCR4 and CCR5. **(A, B)** X4-KO Expi293F cells stably expressing (A) FLAG-CXCR4-VN or (B) FLAG-CCR5-VN were transfected with SSM libraries of (A) myc-CXCR4-VC or (B) myc-CCR5-VC. After gating the main population of viable cells based on forward-side scatter, cells were gated for surface expression of the VC-fused chemokine receptor mutants by detection of the myc tag (expression gate in blue). Within the expression gate, the top 5% of cells for Venus/YFP fluorescence were gated (high BiFC gate in magenta). Cells in the expression gate or in the high BiFC gate were collected in independent sorting experiments.

**Figure 3.**
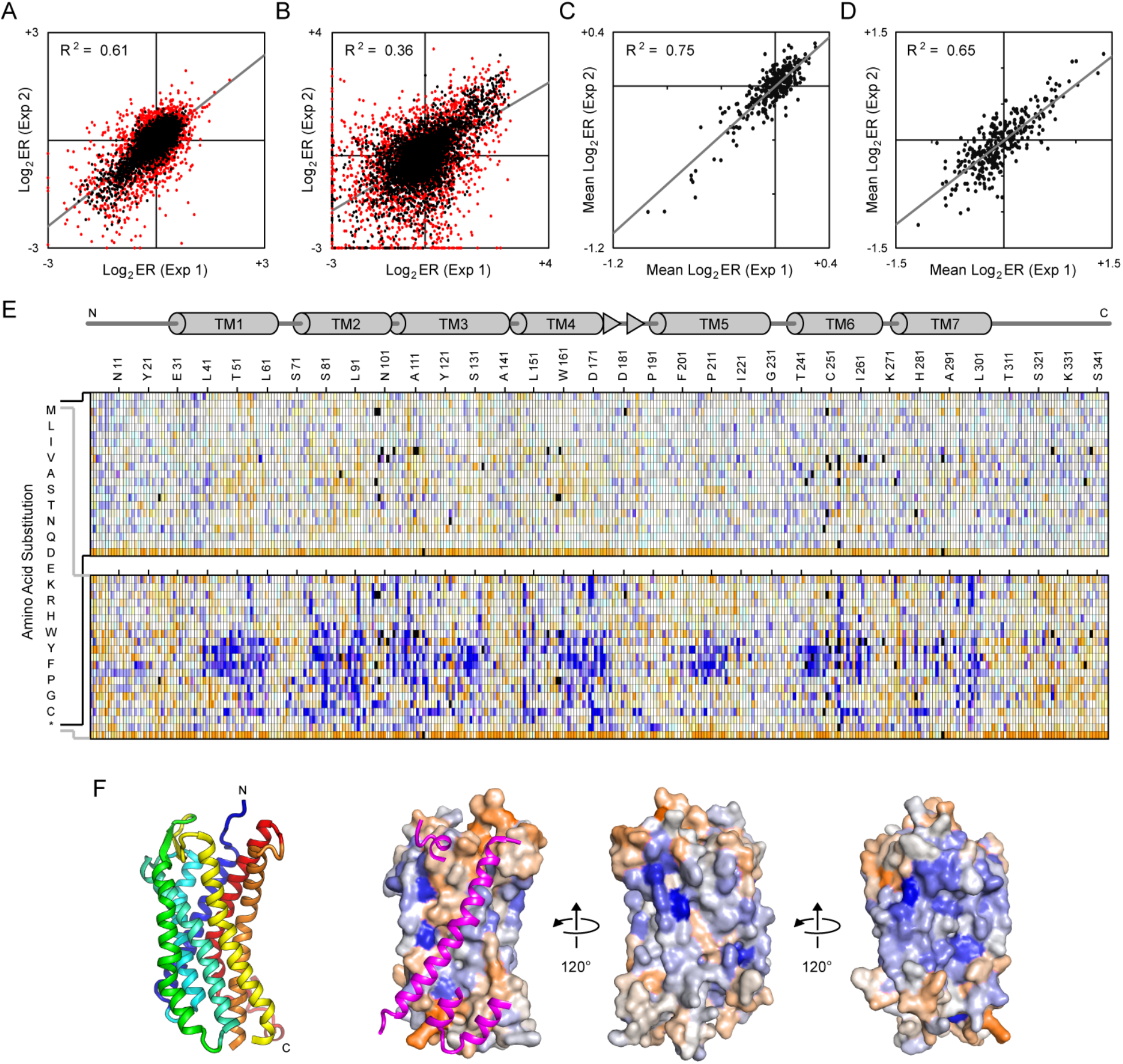
Deep mutational scan and mutational landscape of CXCR4 based on receptor associations. **(A-B)** A single site saturation mutagenesis (SSM) library of CXCR4-VC was expressed in CXCR4-VN Expi293F cells and sorted by FACS for CXCR4-VC surface expression and BiFC. Log2 enrichment ratios for mutations were calculated following Illumina sequencing of the naïve and sorted libraries. Plots show agreement between mutation log2 enrichment ratios after sorting for **(A)** expression or **(B)** expression and BiFC. R-squared values correspond to mutations with frequency ≥ 5×10^−5^ in the naïve library (shown in black). Rare mutations with frequency < 5×10^−5^ in the naïve library are red. Mutations with frequency < 5×10^−6^ were considered absent from the library. **(C-D)** Log_2_ enrichment ratios for all amino acid substitutions at a given residue position were averaged to determine a mean conservation score. Plots show agreement of residue conservation scores between two independent sorting experiments for **(C)** expression or **(D)** expression and BiFC. (**E**) Mutational landscapes of CXCR4-VC sorted for surface expression (top) and BiFC (bottom). Log_2_ enrichment ratios are plotted from depleted/deleterious (≤ -3, orange) to enriched (≥ +3, dark blue). Mutations missing in the naïve library are in black. The CXCR4 sequence is on the horizontal axis and amino acid substitutions are on the vertical axis. A schematic of CXCR4 secondary structure is shown at top, with cylinders representing α-helices and arrows representing β-strands. (**F**) Ribbon structure of CXCR4 (PDB 3ODU) is shown at left, viewed from the plane of the membrane and colored blue to red from N- to C-terminus. The helices forming the dimer interface are facing out. Adjacent on the right, the same orientation of CXCR4 is shown as a surface colored by conservation scores from the BiFC sorting experiment, with conserved residues in orange and residues that are hot spots for enriched mutations in blue. A second CXCR4 molecule is shown as magenta ribbons; for clarity, only regions of the second CXCR4 that are at the dimer interface are shown. To the right, the structure is rotated.

**Figure 4.**
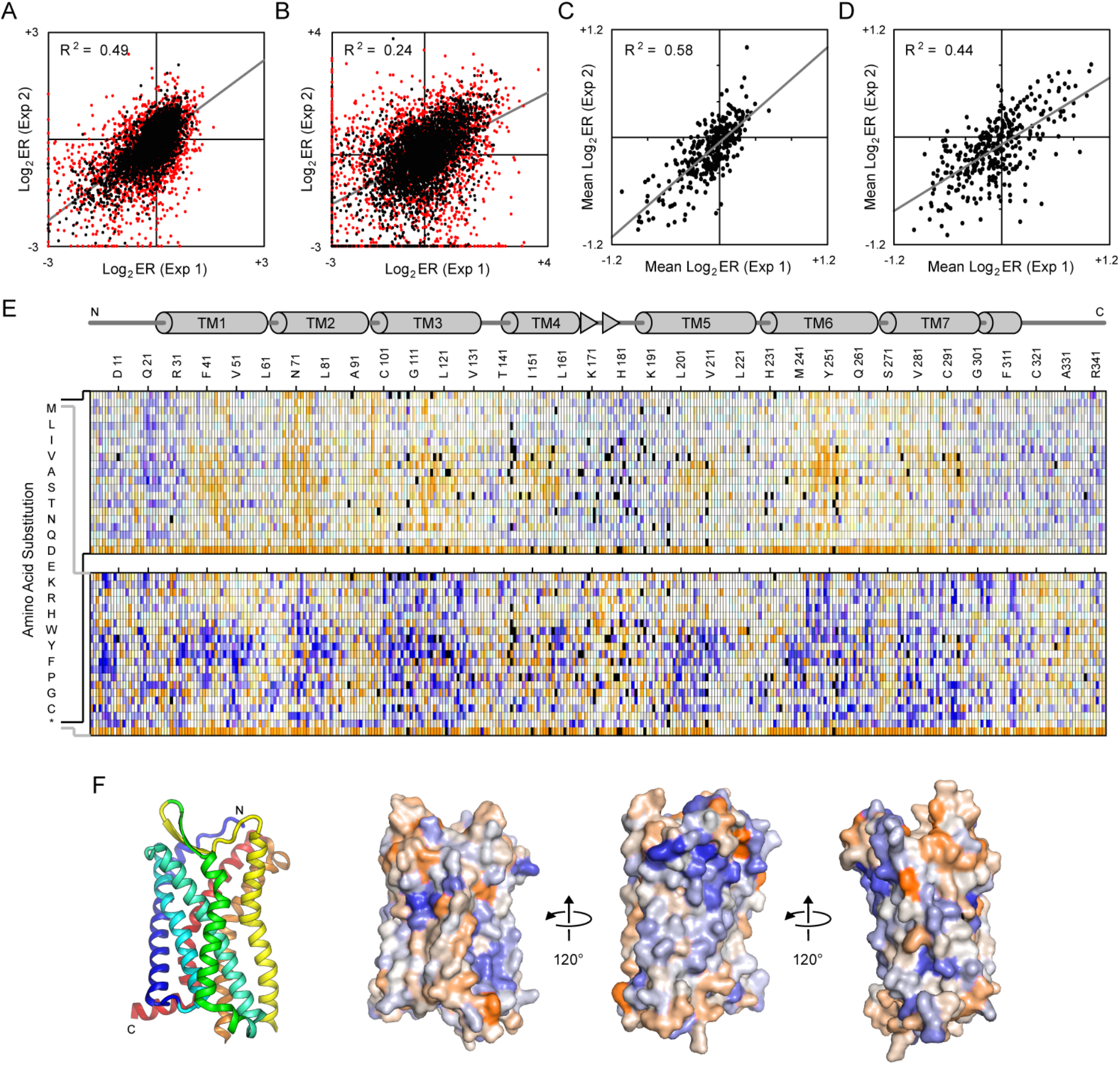
Deep mutational scan and mutational landscape of CCR5 based on receptor associations. **(A-B)** A SSM library of CCR5-VC was expressed in CCR5-VN Expi293F cells and sorted by FACS for CCR5-VC surface expression and BiFC. Agreement between log_2_ enrichment ratios for all mutations is shown when the libraries were sorted for **(A)** expression or **(B)** expression and BiFC. R-squared values correspond to mutations with frequency β 5×10^−5^ in the naïve library (black). Rare mutations with frequency < 5×10^−5^ in the naïve library are red. Mutations with frequency < 5×10^−6^ were considered absent from the library. **(C-D)** Agreement between residue conservation scores from two independent sorting experiments for **(C)** expression or **(D)** expression and BiFC. (**E**) Mutational landscapes of CCR5-VC sorted for surface expression (top) and BiFC (bottom). Log_2_ enrichment ratios are plotted from depleted/deleterious (≤ -3, orange) to enriched (β +3, dark blue). Missing mutations are in black. The CCR5 sequence is on the horizontal axis and amino acid substitutions are on the vertical axis. A schematic of CCR5 secondary structure is shown at top, with cylinders representing α-helices and arrows representing β-strands. (**F**) Ribbon structure of CCR5 (PDB 4MBS) is shown at left, viewed from the plane of the membrane and colored blue to red from N- to C-terminus. The proposed dimer interface of CCR5 is facing out. Adjacent on the right, the same orientation of CCR5 is shown as a surface colored by conservation scores from the BiFC sorting experiment, with conserved residues in orange and residues that are hot spots for enriched mutations in blue. To the right, the structure is rotated.

Sorting experiments were independently replicated. Enrichment ratios weakly agree between the replicate experiments and the agreement is higher when the libraries were sorted for expression only (**Figures 3A and 4A**) versus sorting on expression and BiFC signal (**Figures 3B and 4B**). Low frequency mutations in the naïve library had weaker agreement between the replicate experiments, suggesting low frequency mutations were not deeply sampled during sorting and thus yielded higher variation between replicates. Overall, we consider the ‘noise’ in the data to be similar to earlier deep mutational scans of chemokine receptors (Heredia et al., 2018) but considerably higher than in recent mutational scans of the sweet taste receptor, SARS-CoV-2 receptor ACE2, and serotonin transporter (Chan et al., 2020; Park et al., 2019; Young et al., 2021), which all use equivalent methods. The high noise in this data set may represent difficulties in achieving consistent sort conditions on BiFC signal. Large variations in mutation frequencies in the naïve library is also expected to have contributed to uneven sampling of mutations during sorting of the transfected cell culture. The enrichment ratios for individual mutations should thus be considered as estimates or predictions until verified by targeted mutagenesis. The log_2_ enrichment ratios for all substitutions at a given residue position may also be averaged to calculate a conservation score. Conservation scores show higher agreement between replicate sorting experiments and are considered a more reliable indicator of whether a particular residue is tolerant of mutations (**Figures 3C-D and 4C-D**).

There are two key features in the mutational landscapes. First, for CCR5 but less prominently for CXCR4, mutations of transmembrane residues to polar amino acids tend to reduce expression at the cell surface (**Figures 3E and 4E**). Such mutations are expected to adversely impact folding of the transmembrane domain. This is consistent with previous deep mutational scans of both receptors (Heredia et al., 2018). Second, there are a large number of mutations that are highly enriched for BiFC and these mutations are often found within the transmembrane regions (**Figures 3E and 4E**), where many are expected to be highly destabilizing for tertiary structure (for example, introduction of charged and polar residues in transmembrane helices). It is likely that destabilizing mutations enhance BiFC through non-specific aggregation of mutant proteins. This is consistent with our preliminary data showing that mutations in CCR5 that decreased binding to a conformation-dependent monoclonal antibody (and thus were damaging folded structure) were associated with elevated BiFC. These data indicate the challenges in using mutagenesis to identify dimerization sites in chemokine receptors; a mutation within a dimer interface may disrupt the native dimer but simultaneously increase aggregation and association of misfolded proteins.

To support these hypotheses, we chose 12 mutations for both CXCR4 and CCR5 that were enriched for high BiFC signal. These mutations substitute amino acids for side chains with very different size and chemical properties. In some cases they introduce prolines, which often disrupt secondary structure. We validated that the mutations within CXCR4-VC and CCR5-VC increased BiFC signal in the respective CXCR4-VN and CCR5-VN cell lines, above the BiFC signal of wild type receptors (**Figure 5A,C**). As a control, the metabotropic glutamate receptor mGluR3, which has no known association with chemokine receptors, produced low BiFC signals in these cell lines. Cells were gated based on surface levels of the different receptors to partially control for potential differences in expression, although we do not exclude the possibility that increased BiFC signal arises from elevated levels of aggregating protein trapped in intracellular compartments. Furthermore, we generated a stable mGluR3-VN expressing cell line that provides a high BiFC signal when transfected with mGluR3-VC. mGluR3 is a class C GPCR that forms disulfide bonded dimers. The BiFC signal is low when the mGluR3-VN line is transfected with wild type CXCR4-VC or CCR5-VC. However, the mutant CXCR4 and CCR5 proteins had very high BiFC signal with mGluR3 (**Figure 5B,D**), again suggesting they are non-specifically associating with other proteins in the membrane due to destabilized structure.

**Figure 5.**
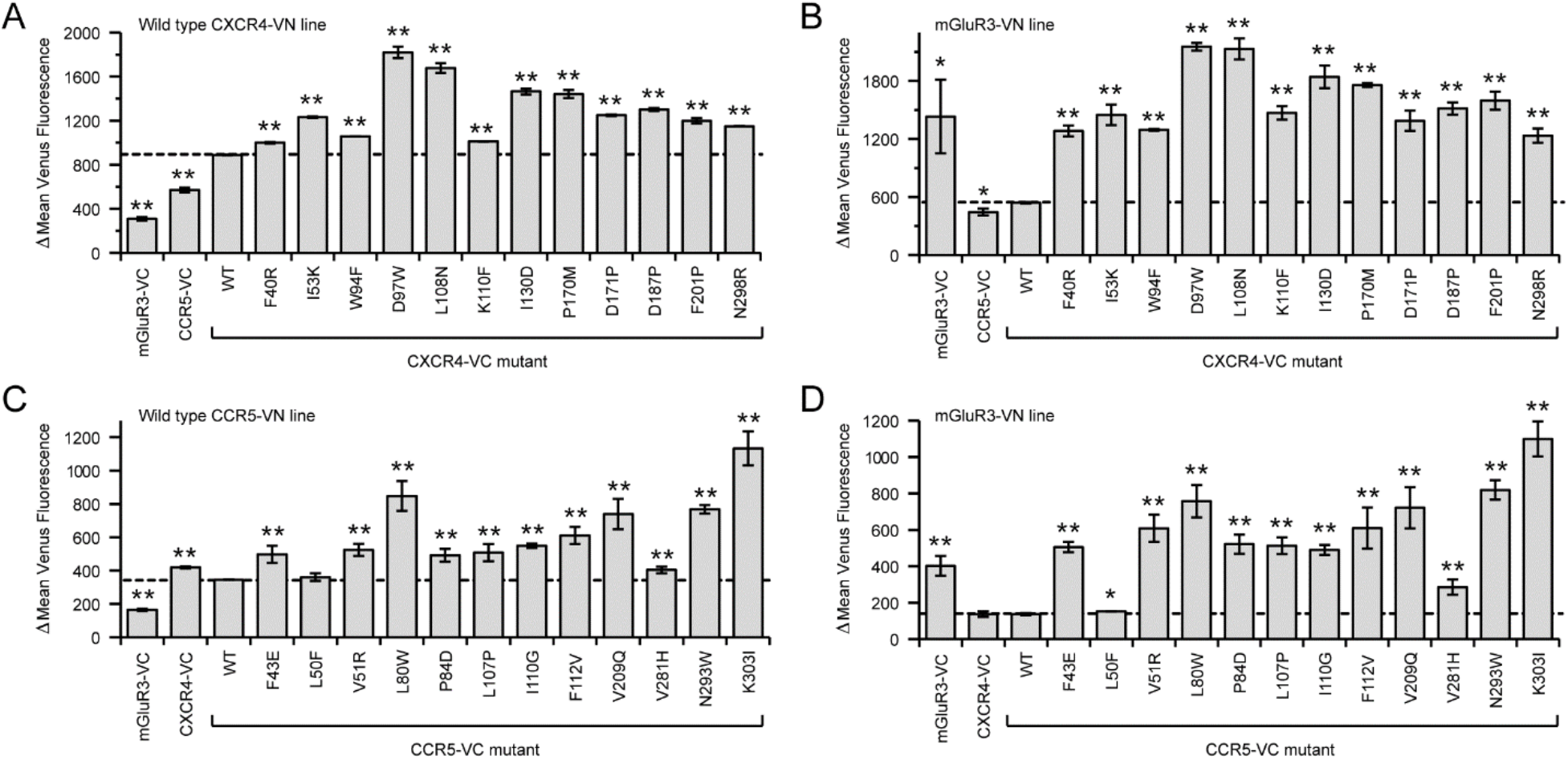
Validation of mutations in chemokine receptors that increase BiFC. **(A,B)** Mutations of CXCR4 predicted from the deep mutational scan to increase BiFC with wild-type CXCR4 were validated. The mutations were introduced into CXCR4-VC and transfected into X4-KO Expi293F cell lines stably expressing either (**A**) CXCR4-VN or (**B**) mGluR3-VN. **(C, D)** Validation of CCR5 mutants enriched in the deep mutational scan for high BiFC signal. The mutations in the CCR5-VC construct were tested in cell lines stably expressing either (**C**) CCR5-VN or (**D**) mGluR3-VN. BiFC signal was measured by flow cytometry. Data are mean ± SD, n=3 independent replicates. *, p < 0.05. **, p < 0.01, Student’s t test.

### Mutations in CXCR4 and CCR5 that reduce receptor associations as measured by BiFC

To understand whether the data provides insights into specific dimer interactions surfaces, the conservation scores from the BiFC selection experiments were mapped to the crystal structures of CXCR4 and CCR5 (**Figures 3F and 4F**). CXCR4 has been crystalized as a dimer, with the same dimeric arrangement being resolved in multiple crystal forms with different molecular packing, providing high confidence that the structure represents a relevant complex (Wu et al., 2010). Residues of CXCR4 known to be buried in the dimeric assembly were weakly more conserved in the BiFC-based selection, with lower mutational tolerance compared to other exposed surfaces of the protein (**Figure 3F**). These more conserved regions are principally on TM3, TM4, and TM5. We note that these regions approximately correspond to the transmembrane surface that was more highly conserved in the published deep mutational scan of CXCR4 for binding to a conformation-dependent antibody (**Figure 1A**). Overall, we conclude that the BiFC-based selection and deep mutational scan succeeded in identifying the known dimer interface of CXCR4. We thus turned our attention to CCR5 where the dimer interface has not been determined by structural methods at atomic resolution. Conservation scores from the BiFC-based deep mutational scan of CCR5 mapped to the protein’s monomeric structure indicate that the transmembrane surface formed by TM3-TM4 has reduced mutational tolerance compared to other surfaces (**Figure 4F**). The TM3-TM4 surface of CCR5 was also more conserved in the published deep mutational scan of CCR5 for binding to a conformation-dependent antibody (**Figure 1**). Accordingly, we hypothesize that this surface is buried in the CCR5 dimer.

We further inspected the mutational landscapes for mutations that were highly depleted in the BiFC-based selection and found a small number in the C-terminal cytoplasmic tails of the receptors. GPCRs are known to cluster in lipid microdomains (Goddard & Watts, 2012; Villar et al., 2016), and their clustering would be anticipated to increase BiFC signal. CXCR4 has a number of basic residues at its C-terminus and CCR5 has palmitoylation sites (Blanpain et al., 2001), both of which are often associated with lipid microdomain localization.

We selected mutations of CXCR4 and CCR5 that were highly depleted following the BiFC-based selection but were close to neutral in the selection for surface expression; low BiFC signal was therefore not predicted to be due to decreased expression. We validated 5-6 mutations in CXCR4-VN and CCR5-VN as decreasing BiFC in the wild-type CXCR4-VC and CCR5-VC stable cell lines (**Figure 6B,E**). We considered that decreased BiFC between a mutant and wild-type receptor might occur if the mutant or wild-type proteins preferentially self-associated, thus excluding VN- and VC-fused proteins from a dimer complex. We therefore introduced the mutations into VN and VC receptors, which were co-transfected **(Figure 6C,F)**. In this arrangement, both receptor chains carry the same mutation. This led to the confirmation of 4 mutants in CXCR4 and 2 in CCR5 as decreasing self-association of receptor chains (**Figure 6H**). In CXCR4, the mutations are P42W, which induces a kink in TM1 and likely has structural consequences; I204W, which is located on TM5 in the crystallographic dimer interface and is thus predicted to disrupt CXCR4 dimers (**Figure 6A**); and R322M and K327W, which change properties in the basic C-terminal tail of CXCR4 and are hypothesized to alter receptor clustering into lipid microdomains. In CCR5, the mutations are V157I, which is exposed to the membrane on TM4 (**Figure 6D**) and is predicted to lie within a dimer interface; and C323I, which modifies a palmitoylation site in the C-terminal tail and is predicted to alter clustering into lipid microdomains.

**Figure 6.**
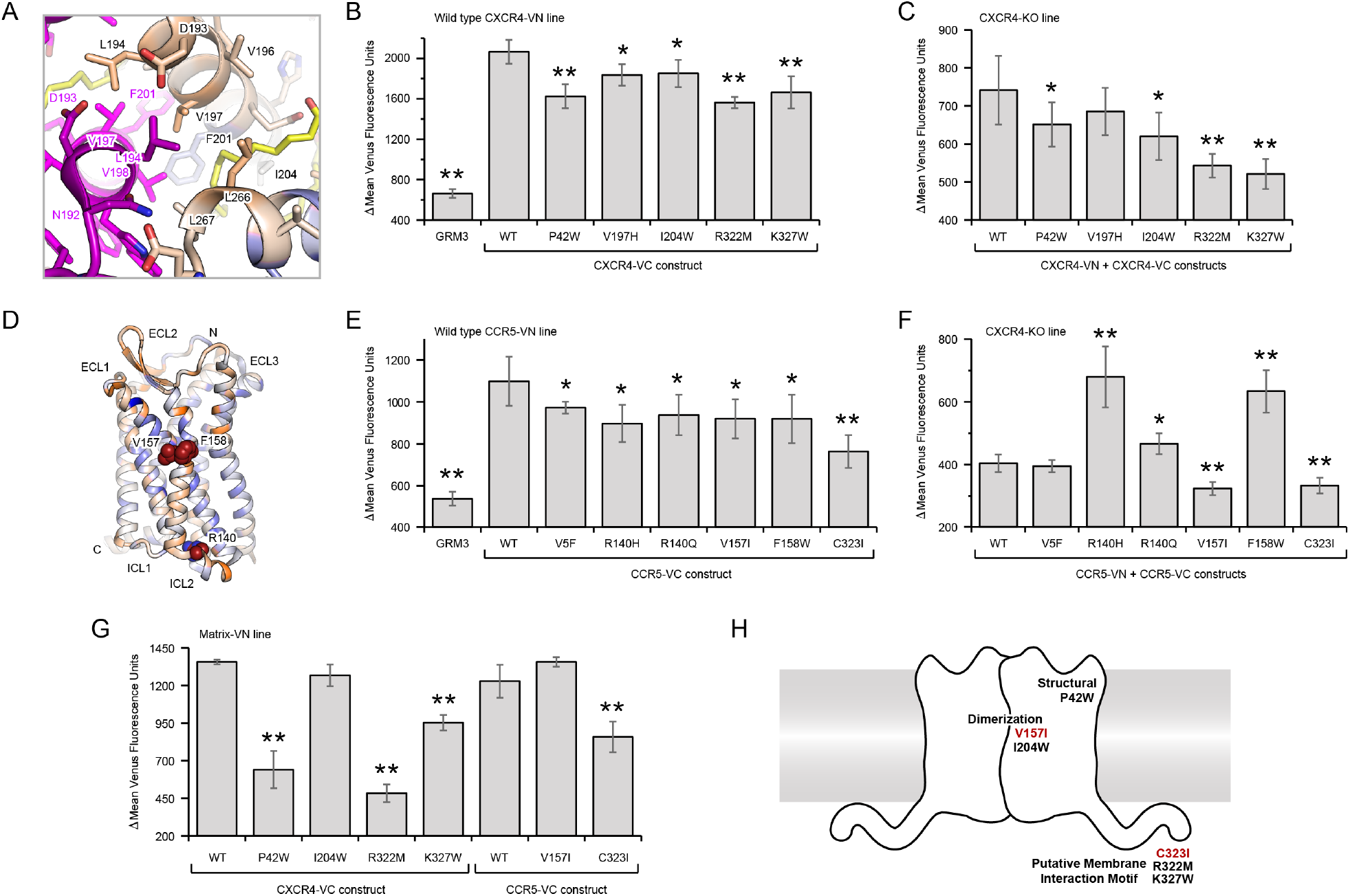
Mutations in CXCR4 and CCR5 that decrease receptor associations based on BiFC signal. **(A)** Dimerization interface of CXCR4 (PDB 3ODU) showing one monomer in magenta and the other monomer colored by the averaged conservation scores from the BiFC-expression deep mutational scan (low mutational tolerance in orange to high mutational tolerance in blue). V197 and I204 are packed against a lipid (yellow sticks) near the interface. **(B, C)** Mutations in CXCR4-VC that were depleted in the BiFC-based deep mutational scan were validated in (**B**) a X4-KO Expi293F cell line stably expressing wild-type CXCR4-VN or (**C**) in X4-KO Expi293F cells in which mutant CXCR4-VC and CXCR4-VN constructs were transiently transfected. (**D**) Ribbon representation of CCR5 (PDB 4MBS) with conservation scores from the BiFC-expression mutational scan mapped to the structure. Residues of low mutational tolerance are orange; residues of high mutational tolerance are blue. A membrane-exposed surface patch that is less tolerant of mutations and is the proposed dimerization interface ir oriented towards the reader, with mutations depleted in the mutational scan in red spheres. **(E, F)** Mutations in CCR5-VC that were depleted in the BiFC-based scan were transfected in (**E**) X4-KO Expi293F cells stably expressing wild-type CCR5-VN and BiFC measured by flow cytometry. The mutations were also tested in (**F**) X4-KO Expi293F cells in which both mutant CCR5-VC and CCR5-VN constructs were cotransfected. (**G**) Mutations in CXCR4-VC and CCR5-VC that reduced BiFC in panels C and F were transfected into a stable MA-VN Expi293F line as a marker for lipid microdomains and BiFC was measured. (**H**) Cartoon of a chemokine receptor dimer annotated with mutations that diminish BiFC between homodimers of CXCR4 (black labels) and CCR5 (red labels). In all data panels, plotted are means ± SD, n=4 biological independent replicates. *, p < 0.05; **, p < 0.01, Student’s t test.

### Effects of CXCR4 and CCR5 mutations

To explore the mechanisms by which these mutations reduce receptor associations, we tested their colocalization with the Matrix protein (MA) of HIV-1, which localizes to cholesterol-rich lipid microdomains (Holm et al., 2003). Expi293F cells that stably express MA fused at the C-terminus to VN were transfected with VC-fused CXCR4 and CCR5 mutants (**Figure 6G**). We found that mutations in the C-terminal tails of CXCR4 (R322M and K327W) and CCR5 (C323I) substantially reduced BiFC interactions with MA, consistent with these mutations reducing localization into lipid microdomains. By comparison, mutations at the putative dimer interfaces (I204W for CXCR4 and V157I for CCR5) had no impact on clustering with MA. Mutation P42W in CXCR4, which is anticipated to have substantial structural effects by targeting a kink-forming proline in TM1, also reduced association with the MA marker for lipid microdomains, indicating that the C-terminal tails are not solely responsible for membrane microdomain localization.

The dynamics of the CXCR4 mutants in the plasma membrane were analyzed by single-molecule TIRF imaging. Halo-tagged CXCR4 proteins were labeled under conditions that permitted tracking of single molecule fluorescence under TIR field. In addition to testing the aforementioned mutants of CXCR4, we included D97W as a control, a mutation located at the end of TM2 and oriented towards the ligand-binding cavity, which was shown to increase BiFC signal (**Figure 5A**). With the exception of CXCR4-D97W, wild-type and mutant CXCR4 proteins were similar, with most molecules mobile within a confined space and with diffusion coefficients < 0.005 μm^2^/s, although a small population of molecules were more mobile (**Figure 7**). Only CXCR4-K327W single molecules had substantially higher mobility based on increased diffusion coefficients. Molecules of the high-BiFC control, CXCR4-D97W, were nearly all immobile (approximately 80%) with low diffusion coefficients, consistent with our speculation that mutations which increase BiFC between receptors promote non-specific aggregation in the membrane.

**Figure 7.**
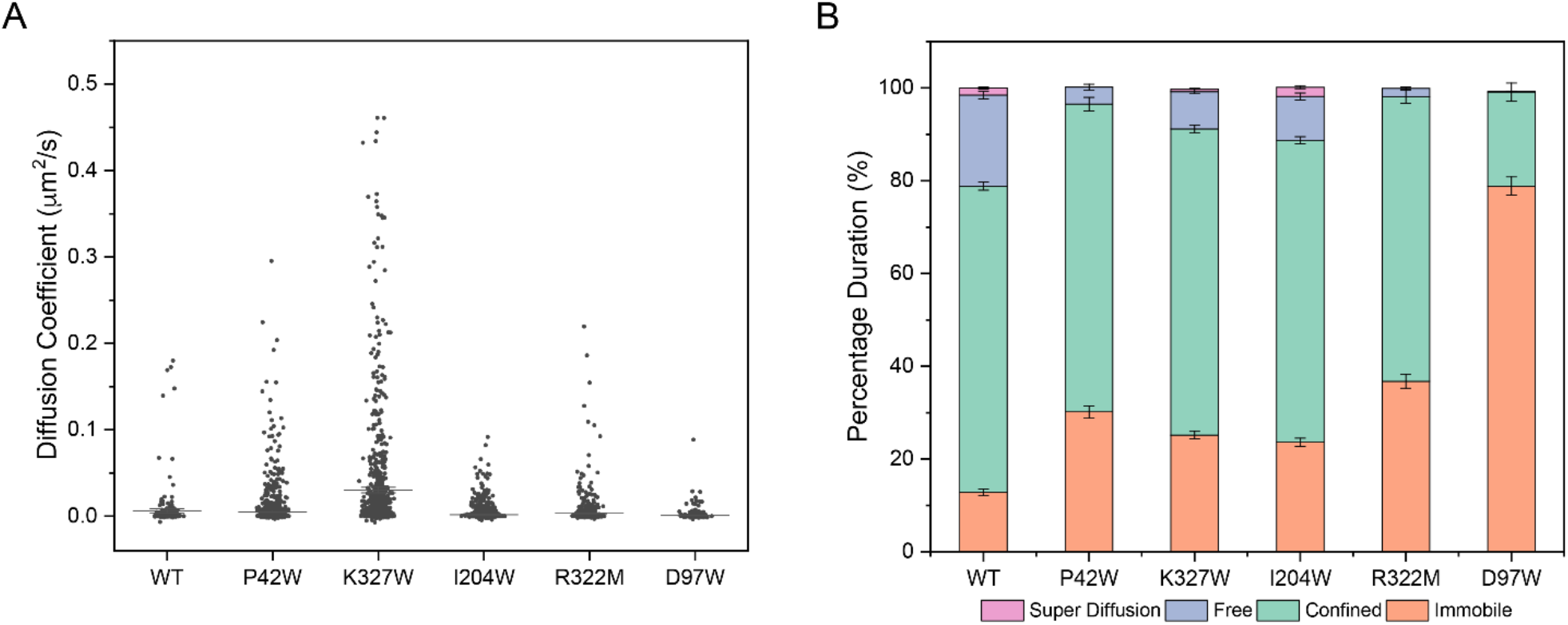
Single molecule imaging of CXCR4. **A)** Single Molecule Imaging analysis of CXCR4 and its mutants. (**A**) Distribution of diffusion coefficients derived from DC-MSS analysis (Vega et al., 2018) for myc-Halo-CXCR4, comparing reduced self-association mutants (P42W, K327W, I204W and R322M) to wild type and one increased self-association mutant (D97W). (**B**) DC-MSS analysis for the tracked trajectories of myc-Halo-CXCR4, dissecting transient motion in the tracks.

The effects of the mutations on ligand binding and calcium mobilization were measured. Using flow cytometry, binding of sfGFP-tagged CXCL12 to the low-BiFC CXCR4 mutants (P42W, I204W, R322M, and K327W) significantly increased compared to wild-type CXCR4 (**Figure 8A**), despite no change or slightly reduced receptor expression at the plasma membrane (**Figure 8B**). CXCL12 binding to CXCR4-P42W and I204W was particularly prominent, increasing 2.5-fold and 5-fold over wild-type, with smaller increases for the two CXCR4 C-terminal mutants. However, increased ligand binding was not associated with higher signaling (**Figure 8C**). Rather, Ca^2+^ mobilization trended lower in the CXCR4 mutants. The CXCR4-P42W mutant was inactive, consistent with P42 having an important structural role in inducing a kink in TM1. The CXCR4-I204W and R322M mutants had decreased signaling, while CXCR4-K327W was similar to wild-type. Hence while the mutations favor high ligand-binding states of CXCR4, those states are decoupled from increased Ca^2+^ signaling. The CCR5-V157I and C232I mutants were also trending towards decreased Ca^2+^ mobilization relative to wild-type CCR5 when stimulated with CCL5 (**Figure 8D**). The data suggest that receptor associations and/or localized clustering, either via oligomeric assemblies or through localization to lipid microdomains, facilitate signaling downstream of ligand binding.

**Figure 8.**
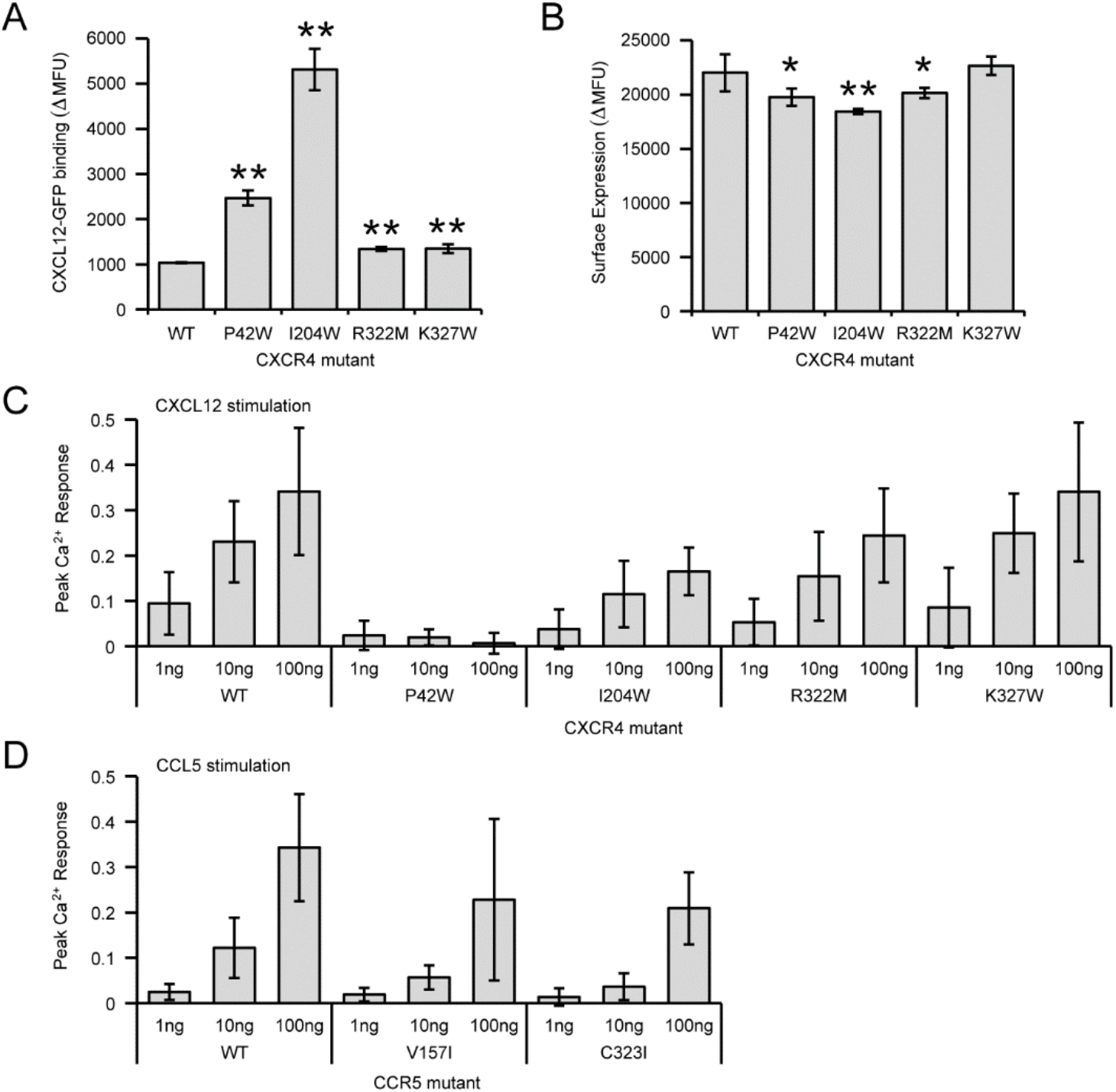
Ligand binding and signaling of chemokine receptor mutants with reduced self-association. **(A)** X4-KO Expi293F cells expressing CXCR4 mutants were incubated with 10 μM CXCL12-sfGFP and bound ligand was detected by flow cytometry. (**B**) Surface expression of CXCR4 mutants measured by flow cytometry. In panels A and B, data are mean ± SD, n=6 biological independent replicates. *, p < 0.05; **, p < 0.01, Student’s t test. **(C)** Calcium mobilization was recorded in Fluo-4-AM loaded X4-KO Expi293F cells transfected with the indicated CXCR4 mutants. Calcium mobilization at 1, 10, and 100 ng/ml CXCL12 concentrations are reported relative to maximum Ca^2+^ response to 4 μM ionomycin. Data are mean ± SD, n=8 biological independent replicates. **(D)** As in panel C, with cells now expressing CCR5 proteins and stimulated with CCL5. Data are mean ± SD, n=6 biological independent replicates.

CXCR4 and CCR5 are coreceptors with CD4 for HIV-1 entry into a host cell. The viral spike protein Env engages CD4 on the host cell, facilitating conformational changes that expose binding sites for CXCR4 (X4-tropic viruses) or CCR5 (R5-tropic viruses) (Chen, 2019). We tested whether the low-BiFC receptor mutations altered syncytia formation when Env-expressing cells were cultured with receptor-expressing cells. Cells expressing CD4 together with wild-type or mutant CXCR4 had no significant differences in syncytia formation when incubated with cells expressing Env from the X4-tropic MN strain (**Figure 9A**), and likewise cells expressing CD4 together with wild-type or mutant CCR5 had no significant differences in syncytia formation when incubated with cells expressing Env from the R5-tropic BaL strain (Dragic et al., 1996) (**Figure 9B**).

**Figure 9.**
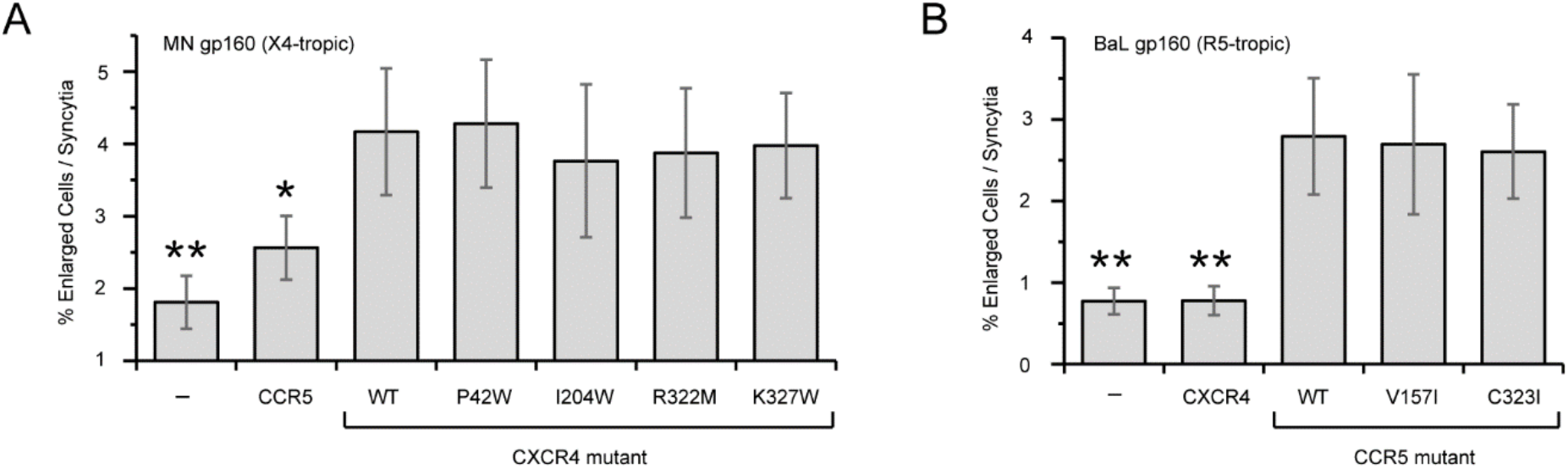
HIV-1 Env-mediated syncytia formation is not affected by mutations in chemokine receptors that reduce self-associations. Formation of syncytia was measured by flow cytometry after co-culturing X4-KO Expi293F cells expressing either (**A**) MN gp160 or (**B**) BaL gp160 with cells expressing CD4 and the indicated CXCR4 or CCR5 mutants, respectively. Data are mean ± SD, n = 6 independent experiments. *, p < 0.05; **, p < 0.01, Student’s t test.

## DISCUSSION

Extensive research efforts have found that chemokine receptors CXCR4 and CCR5 homo- and heterodimerize (Babcock et al., 2003; El-Asmar et al., 2005; Hernanz-Falcón et al., 2004; Issafras et al., 2002; Sohy et al., 2009; Wu et al., 2010), a feature that is common to many other class A GPCRs (Kobayashi et al., 2017; Manglik et al., 2012; Zhao et al., 2019). However, the organization of chemokine receptor dimers and their functional importance remain incompletely understood, in part because identifying mutations that alter dimer strength has been difficult. Here, we set out to define dimer interfaces and find mutations to modulate receptor associations through a BiFC-based deep mutational scan. The use of deep mutagenesis to identify dimerization interfaces was previously applied to the sweet taste receptor and informed the modeling of that receptor’s dimeric architecture, in close agreement with a cryo-EM structure of a related class C GPCR (Park et al., 2019). However, the mutational landscapes we report for CXCR4 and CCR5 unexpectedly revealed many hundreds of mutations that increased receptor associations based on increased BiFC signal. We show that these mutations not only increased BiFC between chemokine receptor chains but also with an unrelated class C GPCR, suggesting the mutations cause non-specific associations and aggregation. This is supported by the observation that the mutations frequently altered chemical properties of amino acids within transmembrane helices and may thus be deleterious for proper folding. Single molecule imaging of CXCR4 with a high-BiFC mutation showed the mutant receptors were mostly immobile in the membrane. The ease with which CCR5 and CXCR4 could be driven towards aggregation means over-expression studies of receptors must be approached with caution, lest non-specific aggregation is falsely interpreted as biologically relevant dimer formation. Despite the high prevalence of BiFC-enhancing mutations in the scans, there were surfaces on CXCR4 and CCR5 where mutations tended to be depleted following BiFC-based sorting. We consider these relatively more conserved surfaces in the sorting experiments to be potential interfaces for dimer formation.

CXCR4 has been crystallized as a symmetric dimer with inter-subunit contacts formed primarily by TM3-5 (Wu et al., 2010). The TM3-5 surface exposed to the membrane is less tolerant of mutations in the BiFC-based deep mutational scan and our data therefore support this dimeric arrangement of CXCR4 in living cells. Substitution I204W in TM5 caused a partial decrease in CXCR4 homodimerization based on BiFC. Cells expressing CXCR4-I204W had 5-fold enhanced binding to CXCL12, consistent with CXCL12/CXCR4 forming a 1:1 complex. However, signaling by the mutant receptor was diminished, although the data do not answer whether decreased signaling was due to reduced dimerization or other structural perturbations caused by the mutation that decouple G protein activation from ligand binding.

The structure of a CCR5 dimer has not been determined experimentally at atomic resolution, although chemical cross-linking has supported close contacts between TM5 helices across the interface (Jin et al., 2018). There is no surface on CCR5 that unequivocally stands out as the putative dimer interface in our BiFC-based deep mutational scan. Our data is most supportive of TM4 forming the CCR5 dimer interface, as the membrane-exposed surface of this helix was relatively more conserved than other transmembrane regions, albeit weakly. It is difficult to compare our findings to the published cross-linking study of CCR5 that emphasized TM5, as cross-linking mutations were not explored in TM4. Substitution V157I, located on TM4 and pointing outwards towards the lipid phase of the membrane, was confirmed as decreasing BiFC between CCR5 receptors.

We also found mutations in the cytoplasmic tails of CXCR4 and CCR5 that decreased BiFC. The cytoplasmic tails are known to contain features that promote lipid microdomain localization (Blanpain et al., 2001; Hancock et al., 1990; Lindwasser & Resh, 2001; Nguyen & Taub, 2002) and decreased BiFC is readily explained by reduced clustering of receptors in membrane microdomains. In agreement, mutations in the C-terminal tails also reduced co-localization with Matrix, a marker for cholesterol-rich microdomains in cellular membranes.

Overall, deep mutational scanning of the chemokine receptors CXCR4 and CCR5 through the use of BiFC demonstrated that these receptors may associate through multiple mechanisms: (1) non-specific associations or aggregation, (2) dimerization, and (3) clustering into lipid microdomains. These findings have implications for studies into chemokine receptor associations, as results that are often interpreted as receptor dimerization or oligomerization may instead arise due to other mechanisms.

## CONFLICTS OF INTEREST

E.P. is an employee and shareholder of Cyrus Biotechnology, Inc. The work described in this manuscript was conducted at the University of Illinois. Cyrus Biotechnology had no role in the project’s design, execution, and analysis/interpretation. The views expressed in the manuscript are solely those of the authors, and they do not represent official views or opinions of the University of Illinois or Cyrus Biotechnology.

## ACKNOWLEDGEMENTS

The UIUC Roy J. Carver Biotechnology Center assisted with deep sequencing and cell sorting. Research was supported by Award Number R01AI129719 from the National Institute of Allergy and Infectious Diseases of the National Institutes of Health to E.P.

